# Human Surfactant Protein D Facilitates SARS-CoV-2 Pseudotype Binding and Entry in DC-SIGN Expressing Cells, and Downregulates Spike protein Induced Inflammation

**DOI:** 10.1101/2022.05.16.491949

**Authors:** Nazar Beirag, Chandan Kumar, Taruna Madan, Mohamed H. Shamji, Roberta Bulla, Daniel Mitchell, Valarmathy Murugaiah, Martin Mayora Neto, Nigel Temperton, Susan Idicula-Thomas, Praveen M Varghese, Uday Kishore

**Affiliations:** Biosciences, College of Health, Medicine and Life Sciences, Brunel University London, Uxbridge UB8 3PH, United Kingdom; Biomedical Informatics Centre, ICMR-National Institute for Research in Reproductive and Child Health, Mumbai 400012, Maharashtra, India; Department of Innate Immunity, ICMR-National Institute for Research in Reproductive and Child Health, Mumbai, India; Immunomodulation and Tolerance Group, Allergy and Clinical Immunology, Department of National Heart and Lung Institute and NIHR Biomedical Research Centre, Asthma UK Centre in Allergic Mechanisms of Asthma, Imperial College London, London, UK; Department of Life Sciences, University of Trieste, Trieste 34127, Italy; WMS - Biomedical Sciences, Warwick Medical School, University of Warwick, Coventry CV4 7AL, United Kingdom; Viral Pseudotype Unit, Medway School of Pharmacy, University of Kent and Greenwich, Kent, ME4 4TB United Kingdom; School of Biosciences and Technology, Vellore Institute of Technology, Vellore, India

**Keywords:** Innate Immune System, Collectins, rfhSP-D, SARS-CoV-2, CoVID-19, Cytokine response

## Abstract

Pattern recognition receptors are crucial for innate anti-viral immunity, including C-type lectin receptors. Two such examples are Lung surfactant protein D (SP-D) and Dendritic cell-specific intercellular adhesion molecules-3 grabbing non-integrin (DC-SIGN) which are soluble and membrane-bound C-type lectin receptors, respectively. SP-D has a crucial immune function in detecting and clearing pulmonary pathogens; DC-SIGN is involved in facilitating dendritic cell interaction as an antigen-presenting cell with naïve T cells to mount an anti-viral immune response. Both SP-D and DC-SIGN have been shown to interact with various viruses, including HIV-1, Influenza A virus and SARS-CoV-2. SARS-CoV-2 is an enveloped RNA virus that causes COVID-19. A recombinant fragment of human SP-D (rfhSP-D) comprising of α-helical neck region, carbohydrate recognition domain, and eight N-terminal Gly-X-Y repeats has been shown to bind SARS-CoV-2 Spike protein and inhibit SARS-CoV-2 replication by preventing viral entry in Vero cells and HEK293T cells expressing ACE2. DC-SIGN has also been shown to act as a cell surface receptor for SARS-CoV-2 independent of ACE2. Since rfhSP-D is known to interact with SARS-CoV-2 Spike protein and DC-SIGN, this study was aimed at investigating the potential of rfhSP-D in modulating SARS-CoV-2 infection. Coincubation of rfhSP-D with Spike protein improved the Spike Protein: DC-SIGN interaction. Molecular dynamic studies revealed that rfhSP-D stabilised the interaction between DC-SIGN and Spike protein. Cell binding analysis with DC-SIGN expressing HEK 293T and THP-1 cells and rfhSP-D treated SARS-CoV-2 Spike pseudotypes confirmed the increased binding. Furthermore, infection assays using the pseudotypes revealed their increased uptake by DC-SIGN expressing cells. The immunomodulatory effect of rfhSP-D on the DC-SIGN: Spike protein interaction on DC-SIGN expressing epithelial and macrophage-like cell lines was also assessed by measuring the mRNA expression of cytokines and chemokines. The RT-qPCR analysis showed that rfhSP-D treatment downregulated the mRNA expression levels of pro-inflammatory cytokines and chemokines such as TNF-α, IFN-α, IL-1β, IL-6, IL-8, and RANTES (as well as NF-κB) in DC-SIGN expressing cells challenged by Spike protein. Furthermore, rfhSP-D treatment was found to downregulate the mRNA levels of MHC class II in DC expressing THP-1 when compared to the untreated controls. We conclude that rfhSP-D helps stabilise the interaction of SARS-CoV-2 Spike protein and DC-SIGN and increases viral uptake by macrophages via DC-SIGN, suggesting an additional role for rfhSP-D in SARS-CoV-2 infection.

## Introduction

Pathogen recognition receptors (PRRs) are germline-encoded host sensors that detect pathogen-associated -molecular patterns (PAMPs) (1). PRRs play a vital part in the regular functioning of the innate immune system (2). They are innate immune system proteins expressed by immune cells, including dendritic cells (DCs), macrophages, neutrophils and monocytes (3). Toll-like receptors (TLRs) and C-type lectin receptors (CLRs) are key of PRRs in host immunity against pathogens (4). Although CLRs are primarily expressed on myeloid cells such as DCs and macrophages, they vary between cell types, allowing specific immune response modifications upon target recognition (5). Receptors such as Dectin-2, Mincle, MGL (Macrophage galactose lectin), Langerin and DC-SIGN (Dendritic Cell-Specific Intercellular adhesion molecules-3-Grabbing Non-integrin) are CLRs that play a major role in the recognition of pathogenic fungi, bacteria, parasites, and viruses (6). The interaction of these CLRs with their ligands allows DCs to moderate the immune response towards either activation or tolerance, which is done through antigen presentation in lymphoid organs and the release of cytokines (7). DCs are responsible mainly for initiating antigen-specific immune responses. Therefore, they are localised at and patrol the sites of first contact with a pathogen, such as mucosal surfaces, including the pulmonary and nasopharyngeal mucosa. Likewise, alveolar macrophages are present in the lung alveoli (8).

DC-SIGN is a CLR that is a surface molecule on DCs that binds to the cell adhesion molecule ICAM-3 on T cells, enhancing DC-T cell contact (9). DC-SIGN is a 44 KDa type II integral membrane protein with a single C-terminal CRD supported by an α-helical neck region with 7 and a half tandem repeats of a 23 amino-acid residue sequence (10, 11). A single transmembrane region anchors the protein, a cytoplasmic domain with recycling, internalisation, and intracellular signalling characteristics(11, 12). DC-SIGN forms oligomers on the cell surface, which improves the avidity of ligand binding and the specificity of binding to multiple repeated units that are likely to be related to the microbial surface features (13). Recently, DC-SIGN has been associated with promoting cis/trans infection of several viruses such as HIV, Cytomegalovirus, Dengue, Ebola and Zika (14-18). The ability of DCs to transmit HIV-1 to CD4^+^ lymphocytes via DC-SIGN coupled with normal DC trafficking suggests that binding of the virus to DC-SIGN could be important in mucosal transmission of HIV-1 because DC-SIGN^+^ DCs are present in the lamina propria at the mucosal surfaces (19). Recently, DC-SIGN has been reported to bind and enhance Severe acute respiratory syndrome coronavirus (SARS-CoV) and SARS-CoV-2 infection independent of ACE2 expression (20).

Another CLRs molecule is human surfactant protein D (SP-D). SP-D belongs to the collectin family with a crucial role in pulmonary surfactant homeostasis and mucosal immunity. SP-D is primarily synthesised and secreted into the air space of the lungs by alveolar type II and Clara cells. Its primary structure is organised into four regions: a cysteine-rich N-terminus, a triple-helical collagen region, and a C-terminal C-type lectin or carbohydrate recognition domain (CRD). SP-D binds to glycosylated ligands on pathogens and initiates opsonisation, aggregation, and direct killing of microbes, facilitating their clearance by phagocytic cells such as macrophages. SP-D was also recently found to bind to the Spike protein of SARS-CoV-2, and inhibit viral replication in Caco-2 cells by promoting viral aggregation, in vitro (21). A recombinant fragment of human SP-D (rfhSP-D), composed of homotrimer neck, CRD, and eight N-terminal Gly-X-Y regions, has been shown to have comparable immunological activities to native SP-D (22). It was shown to bind the HA protein of IAV and act as an entry inhibitor of IAV infection on A549 lung epithelial cells (32). Furthermore, rfhSP-D binds to gp120 and inhibits HIV-1 infectivity and replication in U937 monocytic cells, Jurkat T cells and PBMCs, inhibiting HIV-1 triggered cytokines storm (33). Importantly, rfhSP-D can directly bind to DC-SIGN. This interaction modulates HIV-1 capture and transfer to CD4^+^ T cells (23). Recently, it has been shown rfhSP-D acts as an entry inhibitor of SARS-CoV-2 infection in Vero cells and HEK293T cells expressing ACE2 and TMPRSS2 (24, 25).

SARS-CoV-2, the causative pathogen of Coronavirus Disease 2019 (COVID-19), has resulted in around three million mortality worldwide (26, 27). The most common symptoms are fever, fatigue, and dry cough. The virus has been classified into Alpha, Beta, Gamma, Delta and Omicron variants based on mutations (28, 29). Some individuals can develop severe respiratory distress (30). SARS-CoV-2 is an enveloped RNA virus that uses a homotrimeric glycosylated spike (S) protein to interact with host cell receptors and promote fusion upon proteolytic activation (31). The transmembrane protease TMPRSS2 is known to mediate proteolytic cleavage at the S1/S2 and S2 domains. The receptor binding domain (RBD) is released by S1/S2 cleavage for high-affinity interaction with ACE2, whereas the S2 domain is released by S2 cleavage for effective virus fusion with the plasma membrane (32, 33). As a result, the virus is internalised by the host cells, resulting in viral replication. New copies of SARS-CoV 2 are internalised to infect more cells, increasing the viral load in the lungs, exacerbating the pro-inflammatory response, and extending the cellular and epithelial lung damage (34).

The sequence of events around the Spike protein/ACE2 interaction is well established; however, much remains to be unravelled about additional factors facilitating the infection, such as SARS-CoV-2 delivery to the ACE2 receptor (35). Indeed, Spike protein from both SARS-CoV and SARS-CoV-2 have similar affinity for ACE2 but show very different transmission rates (36, 37). The enhanced transmission rate of SARS-CoV-2 relative to SARS-CoV might result from an efficient viral adhesion through host-cell attachment factor, which may promote efficient infection of ACE2^+^ cells (38, 39). In this framework, DC-SIGN and APCs (DCs and alveolar macrophages) can play a role both in viral attachment and immune activation in the lungs (40-42).

Since rfhSP-D has been shown to inhibit SARS-CoV-2 infection and it binds to DC-SIGN (21, 24, 25), and SARS-CoV-2 spike protein binds to DC-SIGN (20), this study was aimed at investigating whether the interaction of rfhSP-D with SARS-CoV-2 and DC-SIGN exerts antiviral and anti-inflammatory activities.

## Materials and Methods

### Cell Culture and Treatments

HEK 293T cells were maintained in growth media (Dulbecco’s Modified Eagle’s Medium (DMEM) with Glutamax (Gibco) supplemented with 10% v/v foetal bovine serum (FBS), 100U/ml penicillin (Gibco), and 100µg/ml streptomycin (Gibco). The cells were cultured at 37°C in the presence of 5% v/v CO_2_ until 70% confluent. HEK 293T cells were transiently transfected with a plasmid expressing human DC-SIGN (HG10200-UT; Sino Biological), using Promega FuGENE™ HD Transfection Reagent (Fisher Scientific). Next day, the cells were washed and cultured in the presence of hygromycin to select DC-SIGN expressing HEK-293T cells (DC HEK) (Thermo Fisher Scientific). Similarly, THP-1 cells were cultured in growth media. THP-1 cells were induced to express DC-SIGN surface molecules by the treatment with PMA (10□ng/mL) in combination with IL-4 (1000□units/mL) and incubated for 72 h (43).

### Expression and purification of a recombinant fragment of human SP-D (rfhSP-D) containing neck and CRD regions

A recombinant fragment of human SP-D (rfhSP-D) was expressed under bacteriophage T7 promoter in *Escherichia coli* BL21 (λDE3) pLysS (Invitrogen), transformed with for plasmid containing cDNA sequences for neck, CRD regions and 8 Gly-X-Y repeats of human SP-D (24). Briefly, a primary inoculum of 25 ml bacterial culture was inoculated into 500 mL of Luria-Bertani (LB) broth containing 100 µg/ml ampicillin and 34 µg/ml chloramphenicol (Sigma-Aldrich), grown to OD_600_ of 0.6. The bacterial culture was then induced with 0.5 mM isopropyl β-D-1-thiogalactopyranoside (IPTG) (Sigma -Aldrich) for 3 hours. The bacterial cell pellet was harvested and resuspended in lysis buffer (50 mM Tris-HCl pH 7.5, 200 mM NaCl, 5 mM EDTA pH 8, 0.1% Triton X–100, 0.1 mM phenyl-methyl-sulfonyl fluoride (PMSF), 50µg/ml lysozyme) and sonicated (ten cycles, 30 seconds each). The sonicate was harvested at 12000 x g for 30 minutes, followed by solubilisation of inclusion bodies in refolding buffer (50 mM Tris-HCl pH 7.5, 100 mM NaCl, 10 mM 2-Mecraptoethanol) containing 8M urea. The solubilised fraction was dialysed stepwise against refolding buffer containing 4 M, 2M, 1 M and 0M urea. The clear dialysate was loaded onto a maltose-agarose column (5ml; Sigma-Aldrich). The bound rfhSP-D was eluted using 50 mM Tris-HCl pH 7.5, 100 mM NaCl and 10 mM EDTA. The eluted fractions were then passed through a polymyxin B column and sodium deoxycholate buffer (Pierce™ High-Capacity Endotoxin Removal Spin Columns, Thermo Fisher) to remove endotoxin. The endotoxin levels were measured using ToxinSensor™ Chromogenic LAL Endotoxin Assay Kit (Genescript). The amount of endotoxin present in the rfhSP-D batches was ∼ 4 pg/µg of rfhSP-D.

### Expression and purification of soluble tetrameric DC-SIGN

The pT5T construct expressing tetrameric form of human DC-SIGN was transformed into *Escherichia coli* BL21 ((λDE3) pLysS. Protein expression was performed using bacterial culture in LB medium containing 50 µg/ml ampicillin at 37°C until OD_600_ reached 0.7. The bacteria culture was induced with 10 mM IPTG (Sigma -Aldrich) and incubated for 3 h at 37°C. Bacterial cells (1 L) were centrifuged at 4,500 x g for 15 min at 4°C. Next, the cell pellet was treated with 22 ml of lysis buffer containing 100 mM Tris-HCl pH 7.5, 0.5 mM NaCl, 2.5 mM EDTA pH 8, 0.5 mM PMSF, and 50µg/ml lysozyme, and left to stir for 1 h at 4°C. Cells were then sonicated for 10 cycles for 30 s with 2 min intervals. The sonicated suspension was spun at 10,000 g for 15 minutes at 4°C. The inclusion bodies present in the pellet were solubilised in 20 ml buffer containing 10 mM Tris-HCl, pH 7.0, 0.01% β-mercaptoethanol and 6 M urea by rotating on a shaker for 1 h at 4°C. The mixture was then centrifuged at 13,000 x g for 30 min at 4°C. The supernatant was drop-wise diluted fivefold with loading buffer containing 25 mM Tris-HCl pH 7.8, 1 M NaCl and 2.5mM CaCl_2_ with gentle stirring. This was then dialysed against 2 L of loading buffer with three buffer changes every 3 h. Following further centrifugation at 13,000 x g for 15 min at 4°C, the supernatant was loaded onto a mannan-agarose column (5 ml; Sigma) pre-equilibrated with loading buffer. The column was washed with five-bed volumes of the loading buffer, and the bound protein was eluted in 1 ml fractions using the elution buffer containing 25 mM Tris-HCl pH 7.8, 1 M NaCl, and 2.5 mM EDTA. The absorbance was read at 280 nm, and the peak fractions were frozen at -20°C. The purity of the protein was analysed by 15% w/v SDS-PAGE.

### ELISA

Decreasing concentrations of recombinant DC-SIGN or rfhSP-D (2, 1, 0.5 or 0 µg 100µl/well) were coated on polystyrene microtiter plates (Sigma-Aldrich) at 4°C overnight using carbonate/bicarbonate (CBC) buffer, pH 9.6 (Sigma-Aldrich). The microtiter wells were washed three times the next day with PBST Buffer (PBS + 0.05% Tween 20) (Fisher Scientific). The wells were then blocked using 2% w/v BSA in PBS (Fisher Scientific) for 2 h at 37°C and washed three times using PBST. Constant concentration (2 µg 100µl/well) of recombinant SARS-CoV-2 spike protein (RP-87680, Invitrogen) was added to the wells. After a 2-hour incubation at 37°C, the wells were washed with PBST to eliminate any unbound protein. Polyclonal rabbit anti-SARS-CoV-2 spike (NR-52947, Bei-Resources) was used to probe the wells (1:5,000) in PBS and incubated for an additional 1 hour at 37°C. Goat anti-rabbit IgG conjugated to HRP (1:5,000) (Promega) was used to detect the bound protein. The colour was developed using 3,3′,5,5’-tetramethylbenzidine (TMB) (Biolegend), and the reaction was stopped using 1M H_2_SO_4_ (50 µl/well; Sigma-Aldrich) and read at 450 nm spectrophotometrically.

For competitive ELISA, microtiter wells were coated overnight at 4°C with DC-SIGN protein (2 µg; 100 µL /well) and blocked. A fixed concentration of SARS-CoV-2 Spike protein (2µg; 100µl/well) and decreasing concentration (4.0, 2.0, 1.0, 0.5, 0 µg; 100µl/well) of rfhSP-D in calcium buffer, was added to the well as competing proteins. The plate was incubated at 37°C for 1.5 h and then at 4°C for another 1.5 h. To remove any unbound protein, the wells were rinsed three times with PBST. Next, the wells were probed with polyclonal rabbit anti-SARS CoV-2 spike (1:5,000) in PBS and incubated for an additional 1 h at 37°C. The bound protein was detected using goat anti-rabbit IgG conjugated to HRP (1:5000), and the colour was developed using TMB (100 µl/well). The reaction was stopped using 1M H_2_SO_4_ (50 µl/well; Sigma-Aldrich). The plate was read at 450 nm using a microplate reader (BioRad).

### Cell binding assay

SARS-CoV-2 spike pseudotypes used were produced as previously described (44). Briefly, HEK 293Tcells were cultured in growth media to 70-80% confluence at 37°C under 5% v/v CO_2_. Cells were co-transfected using FuGENE® HD Transfection Reagent (Promega) with Opti-MEM® diluted plasmids (450 ng of pCAGGS-SARS-CoV-2 spike, 500ng of p8.91-lentiviral vector and 750 ng of pCSFLW). The transfected cells were incubated for 48h at 37°C under 5% v/v CO_2_. Post incubation, the media containing the pseudotypes was harvested without disturbing the cell monolayer. The media was then passed through a syringe driven 0.45 μm filter to remove any cell debris and the pseudotypes were harvested. The pseudotypes stored at -80°C until further use.

DC-HEK and DC-THP-1 cells were seeded in microtiter wells separately in growth medium (1 × 10^5^ cells/well) and incubated overnight at 37°C. The wells were washed three times with PBS, then rfhSP-D (20 µg/ml), pre-incubated with SARS-CoV-2 spike pseudotypes, were added to the corresponding wells and incubated at room temperature (RT) for 2 h. The microtiter wells were rinsed three times with PBS and fixed with 1% v/v paraformaldehyde (PFA) for 1 min at RT. The wells were washed again with PBS and incubated with polyclonal rabbit anti-SARS-CoV-2 spike (1:200 diluted in PBS) and incubated for 1 h at 37°C. After washing three times with PBST, the corresponding wells were probed with Alexa Fluor 488 conjugated goat anti-rabbit antibody (Abcam) diluted in PBS (1:200) for 1 h at RT. Readings were measured using a Clariostar Plus Microplate Reader (BMG Labtech).

### Fluorescent microscopy

DC-HEK cells were cultured on 13 mm glass coverslips to form a monolayer, followed by incubation with SARS-CoV-2 spike pseudotypes (50µl) at 37°C. For 30 min, cells were rinsed with PBS and fixed using 1% w/v PFA for 1 min. The cells were washed three times with PBS, then blocked with 5% w/v BSA in PBS (Fisher Scientific) for 30 minutes. The cells were incubated for 30-min with mouse anti-human DC-SIGN antibodies to detect DC-SIGN and rabbit anti-SARS-CoV-2 Spike antibodies. Next, cells were washed and incubated with a staining buffer containing Alexa Fluor 647 conjugated goat anti-mouse antibody (Abcam), Alexa fluor 488 conjugated goat anti-rabbit antibody (Abcam), and Hoechst (Invitrogen, Life Technologies). This incubation was done in the dark for 45 min. After rinsing with PBS, the mounted coverslips were visualised on a Leica DM4000 microscope.

### Luciferase reporter activity assay

SARS-CoV-2 spike pseudotypes, pre-incubated with rfhSP-D (20µg/ml), were added to DC-HEK and DC-THP-1 cells separately in a 96-well plate and incubated at 37°C for 24 h. The medium was removed, and cells were washed twice with PBS to remove any unbound SARS-CoV2 spike pseudotypes and rfhSP-D. Fresh growth medium was added and incubated at 37°C for 48h. The cells were washed, and luciferase activity (RLU) was measured using ONE-Glo™ Luciferase Assay System (Promega) and read on Clariostar Plus Microplate Reader (BMG Labtech).

### Quantitative qRT-PCR Analysis

DC-HEK and DC-THP-1 cells (0.5 × 10^6^) were seeded overnight in growth medium. Next day, SARS-CoV-2 Spike protein (500 ng/ml) was pre-incubated with rfhSP-D (20 µg/ml) for 2 h at RT and added to DC-THP-1 cells in serum-free medium. Post incubation at 6h, 12h, 24h and 48h, the cells were washed with PBS gently and pelleted. GenElute Mammalian Total RNA Purification Kit (Sigma-Aldrich) was used to extract the total RNA. After RNA extraction, DNase I (Sigma-Aldrich) treatment was performed to remove any DNA contaminants, then the amount of RNA was quantified at A260 nm using a NanoDrop 2000/2000c (ThermoFisher). The purity of RNA was assessed using the ratio A260/A280. Two micrograms of total RNA were used to synthesise cDNA, using High-Capacity RNA to cDNA Kit (Applied Biosystems). The primer BLAST software (Basic Local Alignment Search Tool) was used to design primer sequences as listed in Table 1. The qRT-PCR assay was performed using the Step One Plus system (Applied Biosciences). Each qPCR reaction was conducted in triplicates, containing 75 nM of forward and reverse primers, 5 µl Power SYBR Green Master Mix (Applied Biosystems), and 500 ng of cDNA. qPCR samples were run for 50 °C, and 95°C for 2 and 10 min, followed by running the amplification template for 40 cycles, each cycle involving 15 s at 95 °C and 1 min at 60°C. 18S rRNA was used as an endogenous control to normalise the gene expression.

**Table 1:**
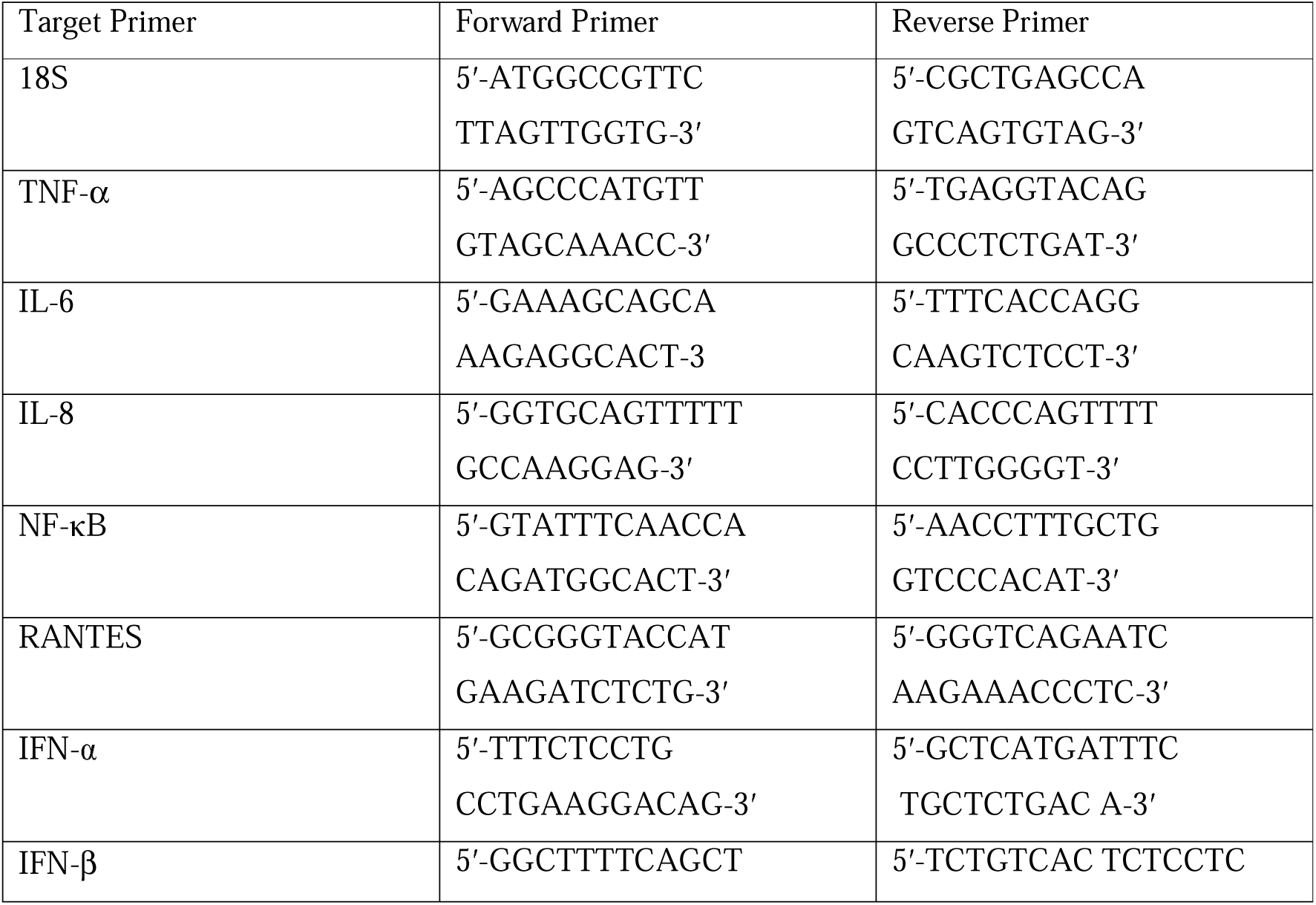
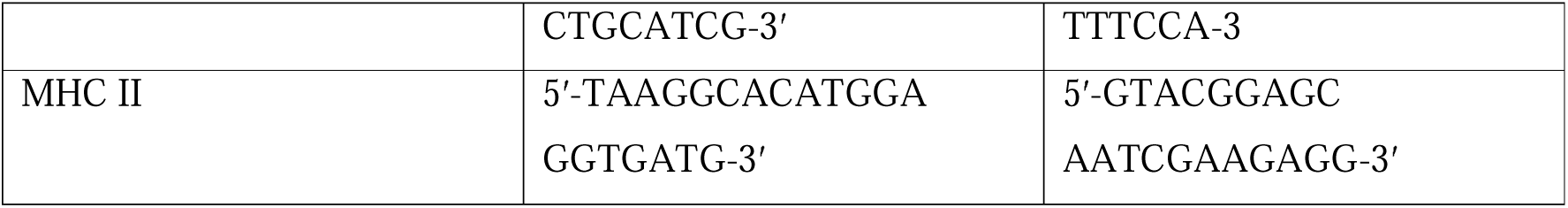
Forward and reverse primers used for RT-qPCR

### Molecular docking

Tripartite complex models of DC-SIGN tetramer, Spike trimer and rfhSP-D trimer were predicted through blind molecular docking using ZDOCK module of Discovery Studio 2021. The structural coordinates for DC-SIGN (CRD), spike and rfhSP-D were retrieved from PDB with IDs as 1K9I, 6XM3, and 1PW9, respectively. Docking was performed in two stages. In the first stage, DC-SIGN (CRD) tetramer was blind docked individually with rfhSP-D trimer (complex A) and spike trimer (complex B). The top ranked poses were analysed for intermolecular interactions and corroborated based on previous studies (23).

In the second stage, the selected docked pose of complex A was further blind docked with spike trimer to build a tripartite complex of DC-SIGN (CRD), Spike and rfhSP-D (complex C). The tripartite complex was selected based on the docking score and intermolecular interactions that were in agreement with literature reports (24).

### Molecular dynamics (MD) simulation

MD simulations for the complexes B, C1 and C2 were performed using GROMACS v2020.6 (44). The force field AMBER99SB was applied with improved protein side-chain torsion potentials (45). All the three complexes were solvated in triclinic periodic box condition using TIP3P water molecules with a distance of 1.5 nm from the center of the complex. Complexes were neutralized by adding Na^+^ counter ions and subsequently minimized for 5000 energy steps using steepest descent algorithm with a tolerance of 1000 kJ/mol/nm. Equilibration was performed using NVT and NPT ensembles for 50,000 steps. Finally, MD was run at constant temperature (300 K) and pressure (1 atm) for 20ns. The analyses of obtained MD trajectories were carried out using GROMACS utility tools.

### Statistical analysis

Graphs were generated using GraphPad Prism 8.0 software. The statistical significance was considered as indicated in the figure legends between treated and untreated conditions. Error bars show SD or SEM as stated in the figure legends.

## Results

### Both DC-SIGN and rfhSP-D Bind to SARS-CoV 2 Spike protein

DC-SIGN and rfhSP-D were expressed in E. coli and purified on mannose and maltose agarose affinity columns, respectively. An indirect ELISA was performed by coating microtiter plates with decreasing concentration with either rfhSP-D or DC-SIGN and probing with anti-SARS-CoV-2 spike antibody to confirm the protein-protein interactions between the two proteins. Both DC-SIGN (Figure 1A) and rfhSP-D (Figure 1B) independently exhibited a dose-dependent increase in binding at all tested concentrations. Since both rfhSP-D and DC-SIGN bound SARS-CoV-2 Spike protein independently, a completive ELISA was performed to evaluate if rfhSP-D would interfere with the binding between Spike protein and DC-SIGN. As a matter of fact, addition of rfhSP-D enhanced the binding of DC-SIGN to the Spike protein in a dose-dependent manner (Figure 1C).

**Figure 1:**
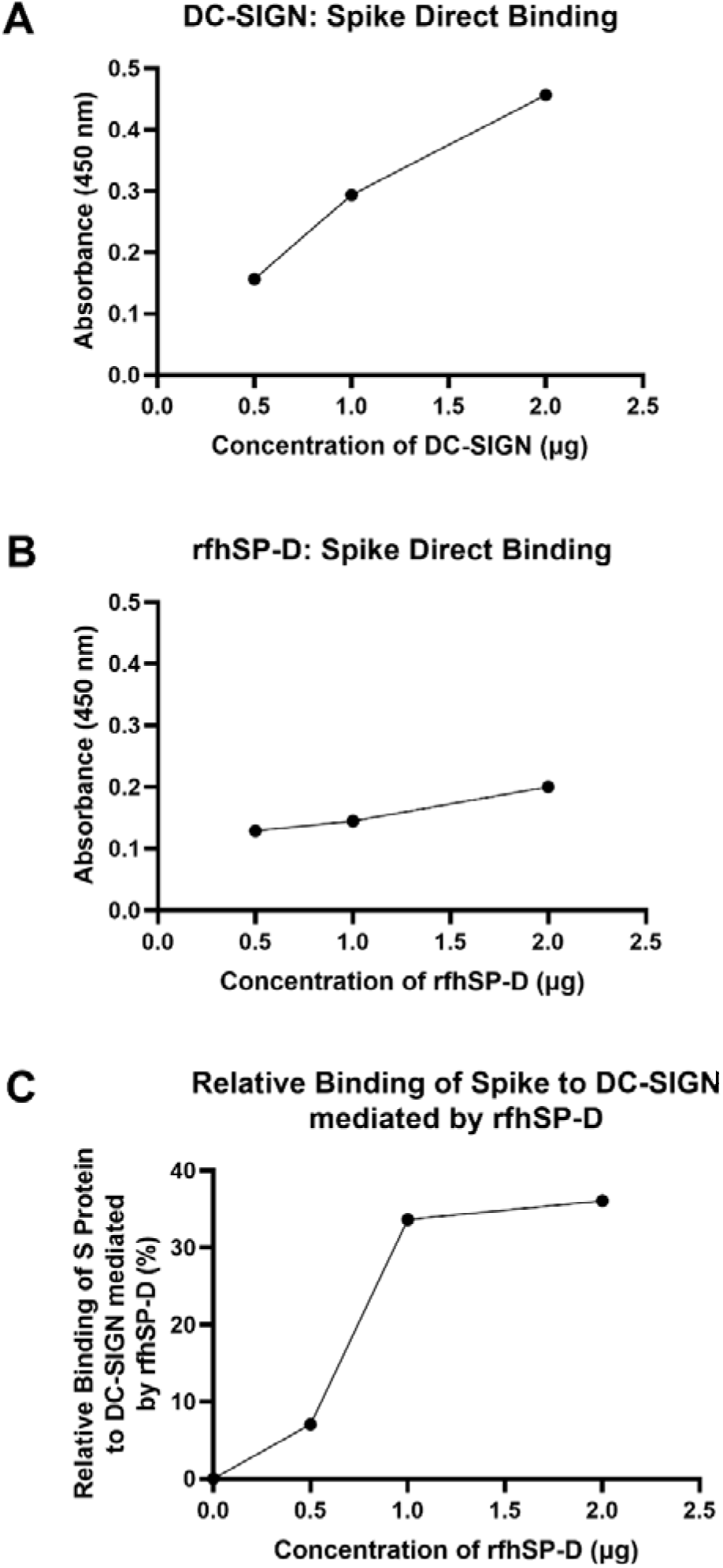
rfhSP-D promotes interaction between SARS-CoV-2 Spike protein and DC-SIGN. The binding of immobilised DC-SIGN (A) or immobilised rfhSP-D (B) to SARS-CoV-2 spike protein was analysed by ELISA. Microtiter wells were coated with a decreasing concentration of DC-SIGN or rfhSP-D (2, 1, 0.5 or 0 µg per well) proteins and incubated with a constant amount of SARS-CoV-2 Spike protein (2 µg per well). Both proteins were found to bind Spike protein in a dose-dependent manner. Competitive ELISA (C) was performed to analyse the effect of rfhSP-D on DC-SIGN: Spike protein interaction. rfhSP-D brought about increased binding between Spike protein and DC-SIGN. Since increasing the concentration of rfhSP-D was found to increase the detectable amount of Spike protein, it seems to suggest the existence of distinct binding sites for the Spike protein on both C type lectins. The data were expressed as a mean of three independent experiments done in triplicates ± SEM.

### rfhSP-D treatment enhances DC-SIGN mediated binding and uptake of SAR-CoV-2 Pseudotyped Viral Particles

Since rfhSP-D was found to interact with DC-SIGN and Spike protein, we evaluated the ability of rfhSP-D to mediate the binding of SARS-CoV-2 to DC-SIGN expressing cells. HEK 293T cells were transfected with a construct containing a DNA sequence of full-length human DC-SIGN to induce DC-SIGN cell surface expression. As previous studies have established the ability of SARS-CoV-2 spike protein to bind DC-SIGN, the binding of the SARS-CoV-2 spike protein-expressing pseudotypes to DC-HEK cells was also confirmed microscopically (Figure 2). To assess the effect of rfhSP-D on pseudotypes binding to DC HEK cells, the cells were challenged with rfhSP-D (20µg/ml) treated SARS-CoV-2 Spike protein-expressing pseudotypes. Increased binding (∼50%) in the treated samples (DC-HEK + SARS-CoV-2 spike Pseudotypes + rfhSP-D) compared to their untreated counterparts (DC-HEK + SARS-CoV 2 spike Pseudotypes) was observed (Figure 3A). The quantitative evaluation of the binding of the pseudotypes was also performed using THP-1 cells treated with PMA and IL-4 to induce the expression of native DC-SIGN. A similar result was obtained using DC-SIGN expressing THP-1 macrophage-like cells. rfhSP-D treatment was found to increase the binding efficiency of the pseudotypes to the THP-1 cells expressing DC-SIGN by ∼ 25%, compared to the untreated controls (Figure 3B).

**Figure 2:**
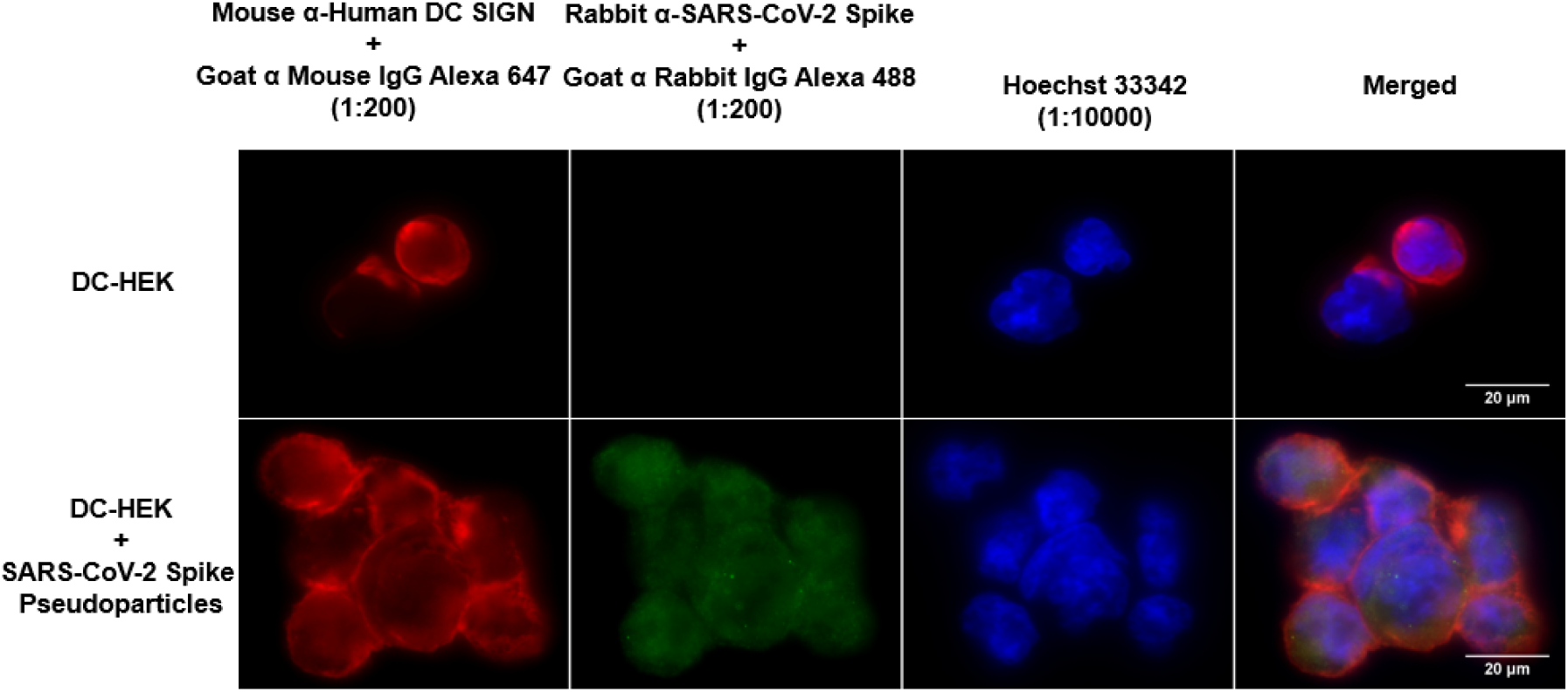
Binding of SARS-CoV-2 Spike-Pseudotypes to DC-SIGN expressing cells. DC-HEK cells were incubated with SARS-CoV-2 Spike Pseudotypes for 30 min at 37°C. Spike Pseudotypes challenged DC-HEK cells were fixed with 4% paraformaldehyde, washed, and blocked with 5% FCS. The cells were probed with rabbit anti-SARS-CoV-2 Spike antibody and mouse anti-DC-SIGN to detect the presence of Spike-Pseudotypes and DC-SIGN expressed on the cells, respectively. Alexa Fluor 647 conjugated goat anti-mouse antibody (Abcam), Alexa fluor 488 conjugated goat anti-rabbit antibody (Abcam), and Hoechst (Invitrogen, Life Technologies) were used to detect the primary antibodies and nucleus.

**Figure 3:**
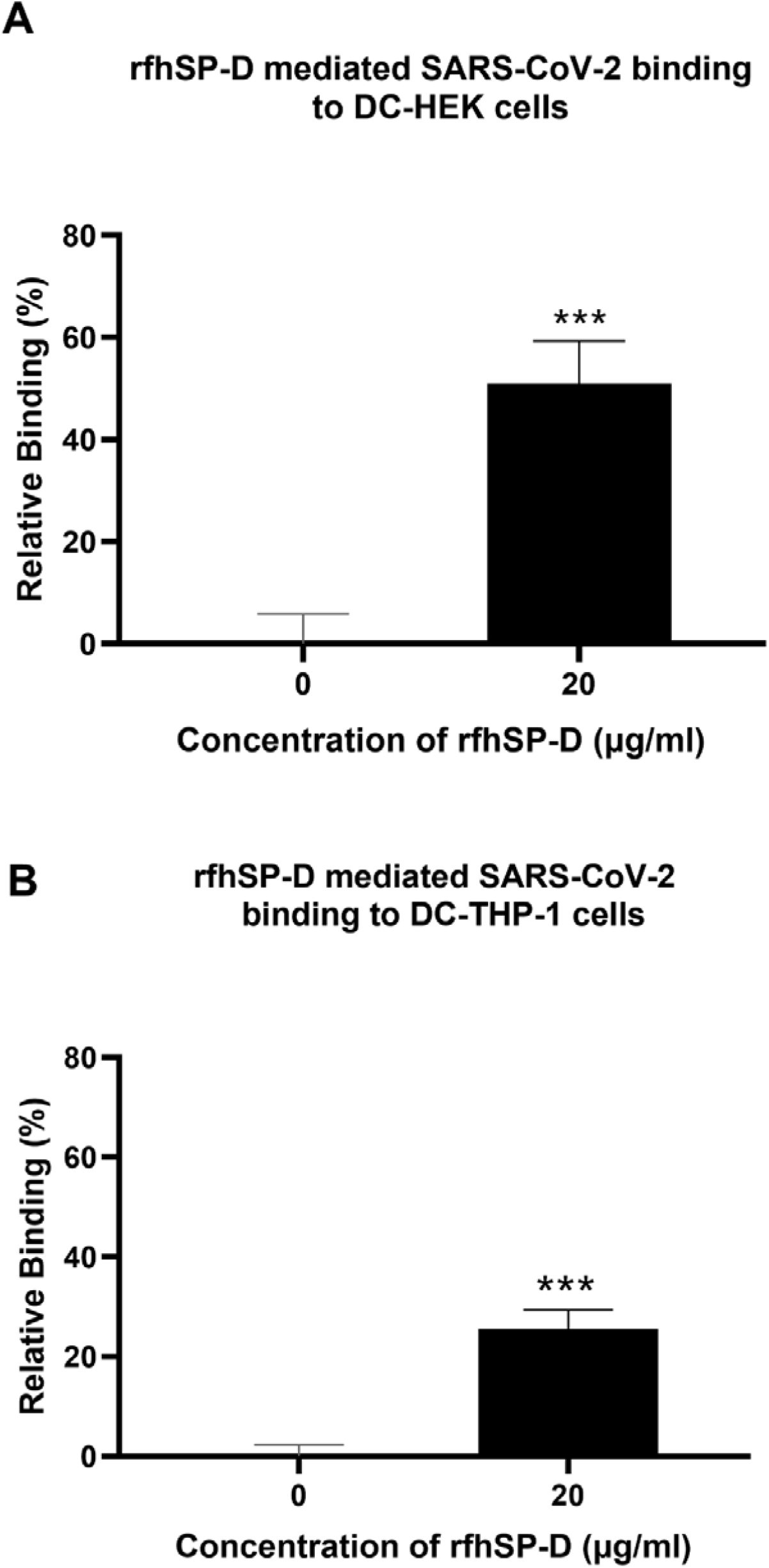
rfhSP-D promotes interaction between SARS-CoV-2 Spike Pseudotypes with DC-SIGN expressing cells. DC-HEK cells (A) and DC-THP-1 cells (B) were treated with rfhSP-D and SARS-CoV-2 Spike-Pseudotypes. The cell binding was analysed using Alexa Fluor 488 (FTIC) and Alexa Fluor 647 (APC); the fluorescence intensity was measured using a GloMax 96 Microplate Luminometer (Promega). An increased fluorescence intensity was observed in DC-HEK and DC-THP-1 cells treated with 20 µg/ml of rfhSP-D compared to cells challenged with Spike pseudotypes alone. Experiments were conducted in triplicates, and error bars represent ± SEM. Unpaired t-test was used calculate the significance (*p < 0.05, **p < 0.01, and ***p < 0.001) (n = 3). (0, untreated sample; 20, treated sample).

To evaluate the impact of rfhSP-D on the transduction of pseudotypes to DC-HEK cells, the cells were treated with rfhSP-D (20µg/ml), challenged with SARS-CoV 2 S protein-expressing pseudotypes for 24h. Higher luciferase activity (∼190 %) in the treated samples (DC-HEK + SARS-CoV-2 spike pseudotypes + rfhSP-D) as compared with their untreated counterparts (DC-HEK + SARS-CoV-2 spike pseudotypes) was noticed (Figure 4A). A similar observation was made using DC-SIGN expressing DC-THP-1 cells compared to untreated controls. rfhSP-D treatment increased the transduction effectiveness of the pseudotypes in DC-THP-1 cells by ∼ 90% (Figure 4B).

**Figure 4:**
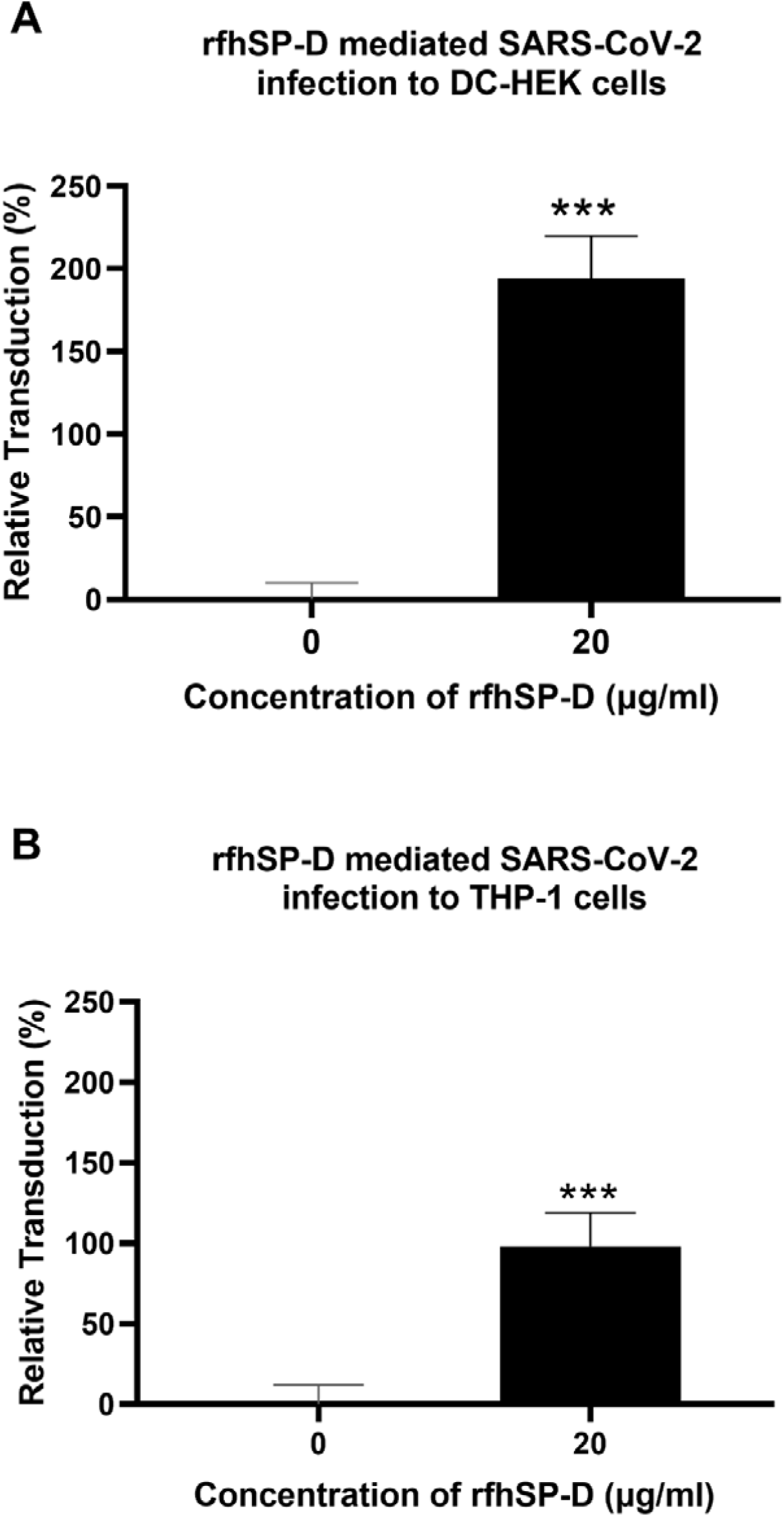
rfhSP-D enhances SARS-CoV-2 Spike pseudotypes transduction by DC-HEK and DC-THP-1 cells. Purified Spike pseudotypes were used to transduce DC-HEK (A) and DC-THP-1 cells (B), and the luciferase reporter activity was measured. Higher levels of luciferase reporter activities were observed in DC-HEK and DC-THP-1 cells when treated with 20 µg/ml of rfhSP-D compared to cells challenged with Spike pseudotypes only. Experiments were conducted in triplicates, and error bars represent ± SEM. Unpaired t-test was used calculate the significance (*p < 0.05, **p < 0.01, and ***p < 0.001) (n = 3).

### rfhSP-D Modulates Pro-Inflammatory Cytokines and Chemokines Response in SARS-CoV-2 Spike protein Challenged DC-HEK cells

Pro-inflammatory cytokines and chemokines such as TNF-α, IFN-α, RANTES, and NF-κB transcription factors characterise SARS-CoV-2 infection in the lower respiratory epithelium that express DC-SIGN. DC HEK cells were challenged with SARS-CoV-2 Spike protein pre-incubated with rfhSP-D to understand better the effect of rfhSP-D on the pro-inflammatory cytokines/chemokines released during SARS-CoV-2 infection. The total RNA extracted from the cells was then used in qRT-PCR, with cells challenged with SARS-CoV-2 Spike protein that had not been treated with rfhSP-D serving as the control. rfhSP-D treatment decreased mRNA levels of TNF-α, IFN-α, RANTES, and NF-κB in DC-HEK cells were challenged with Spike protein. TNF-α mRNA levels were reduced by (∼ -3.3 log_10_) (Figure 5C), while IFN-α, the levels were downregulated (∼ -2.1 log_10_) (Figure 5B). As RANTES response is induced by detection of viral components within infected cells, rfhSP-D treatment reduced the mRNA levels of RANTES in DC-HEK cells challenged with Spike by (∼ -1.3 log_10_). Antiviral cytokines/chemokines are regulated by the transcription factor NF-κB; NF-κB mRNA levels were reduced (∼ -1.2 log_10_) (Figure 5A).

**Figure 5:**
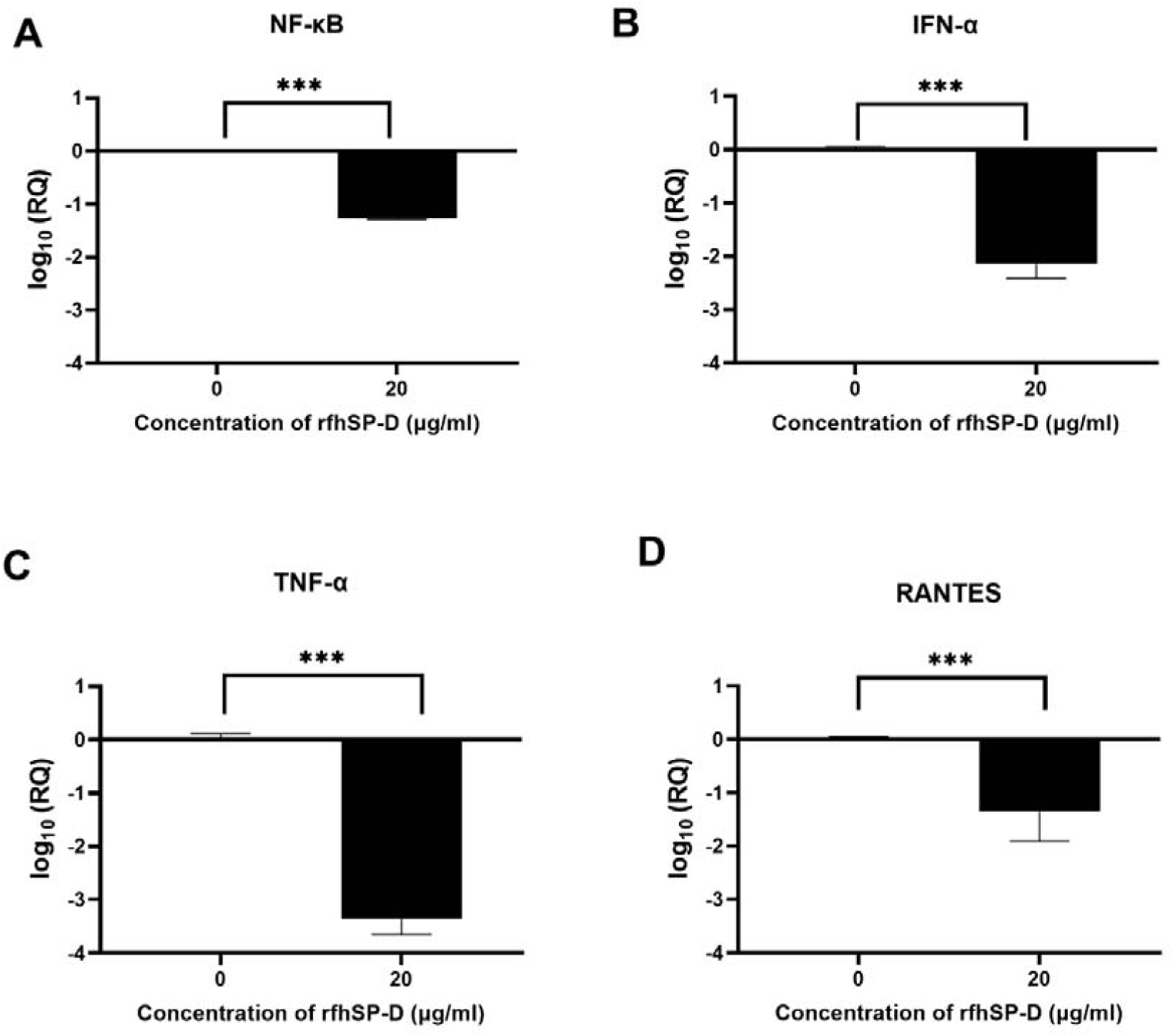
rfhSP-D downregulates pro-inflammatory cytokines and chemokines in DC-HEK cells. SARS-CoV-2 Spike protein incubated with 20μg/ml of rfhSP-D was used to challenge DC-HEK cells. Cells were harvested at 6 h to analyse the expression of cytokines. RNA was purified and converted into cDNA. The gene expression levels of cytokines NF-κB (A), IFN-α (B), TNF-α (C), and RANTES (D) were assessed using RT-qPCR. 18S rRNA was used as an endogenous control. The relative expression (RQ) was calculated using cells challenged with Spike protein untreated with rfhSP-D as the calibrator. The RQ value was calculated using RQ = 2^−ΔΔ^Ct. Assays were conducted in triplicates, and error bars represent ± SEM. Significance was determined using the two-way ANOVA test (**p < 0.01, and ****p < 0.0001) (n = 3).

### Modulation of Immune Response in SARS-CoV-2 Spike Protein-Challenged DC-THP-1 cells by rfhSP-D

Lung macrophages secrete pro-inflammatory mediators such as IL-1, IL-6, IL-8, and TNF-α in response to SARS-CoV-2 infection. To further understand the role of rfhSP-D in producing pro-inflammatory cytokines/chemokines from lung macrophage expressing DC-SIGN during SARS-CoV-2 infection, rfhSP-D treated/untreated SARS-CoV-2 Spike protein was used to challenge DC-THP-1 cells. Following that, qRT-PCR was used to assess the mRNA levels of pro-inflammatory cytokines and chemokines in cells after treatment at 6h and 12h time points (Figure 6). In DC THP-1 cells challenged with Spike protein, rfhSP-D treatment reduced mRNA levels of IL-1, IL-6, IL-8, TNF-α, and NF**-**κB (Figure 6). mRNA levels of NF**-** κB at 6h were slightly reduced (∼ -1 log_10_). At 12 h, it was significantly downregulated (∼ -4 log_10_) in rfhSP-D treated DC-THP-1 cells challenged with Spike protein (Figure 6A). Cells challenged with Spike protein and treated with rfhSP-D at 6h and 12h exhibited a reduction in the gene expression levels of TNF-α (∼ -3.1 log_10_ and ∼ -6.8 log_10_, respectively) (Figure 6B). In rfhSP-D treated DC THP-1 cells challenged with Spike protein, IL-1β mRNA levels were reduced (∼ -2.5 log_10_) after 6 h and (∼ -4 log_10_) 12 h after treatment (Figure 6C). Furthermore, IL-6 levels were significantly downregulated at 12h (-5 log_10_) in rfhSP-D treated DC-THP-1 cells challenged with Spike protein (Figure 6D). Reduced levels of IL-8 at 6h (∼ -2.3 log_10_) and 12h (∼ -4.8 log_10_) were detected in DC-THP-1 cells challenged with Spike protein, treated with rfhSP-D (Figure 6E). MHC class II molecules play a key role in bridging innate immunity to adaptive immunity during anti-viral immune response. rfhSP-D reduced MHC class II expression levels at 6 (-2 log_10_) and 12h (-2.7 log_10_) in DC-THP-1 cells challenged with Spike protein (Figure 6F).

**Figure 6:**
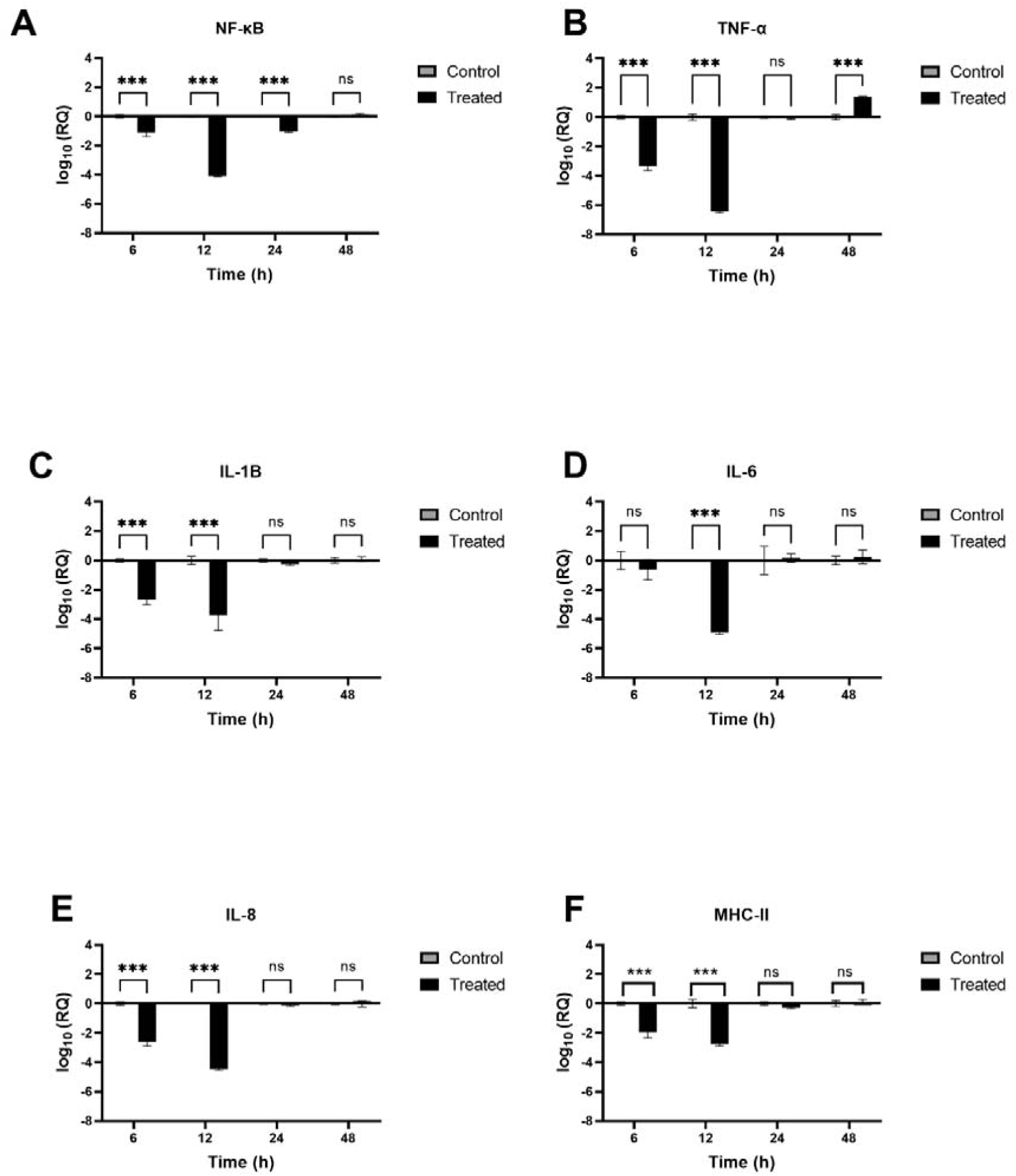
rfhSP-D modulates immune response in DC-THP-1 cells. SARS-CoV-2 Spike protein incubated with 20μg/ml of rfhSP-D was used to challenge DC-THP-1 cells. Cells were harvested at 6h, 12h, 24h, and 48h to analyse the expression of cytokines and MHC class II. Cells were lysed, and purified RNA was converted into cDNA. The expression levels of cytokines NF-κB (A), TNF-α (B), IL-1β (C), IL-6 (D), IL-8 (E) and MHC class II (F) were measured using RT-qPCR, and the data were normalised against 18S rRNA expression as a control. Experiments were conducted in triplicates, and error bars represent ± SEM. The relative expression (RQ) was calculated using cells challenged with Spike protein untreated with rfhSP-D as the calibrator. RQ = 2^− ΔΔ^Ct was used to calculate the RQ value. Significance was determined using the two-way ANOVA test (**p < 0.01, and ****p < 0.0001) (n = 3).

### SP-D interacts with RBD and DC-SIGN interacts with NTD of SARS-CoV-2 Spike protein

DC-SIGN and SP-D are known to interact through their CRDs (23). This interaction was observed in complex A (docked pose 2) of the current study (Figure 7A & table 2). The binding site of DC-SIGN (CRD) and Spike protein is not known; therefore, a blind docking approach was attempted to generate complex B. Analysis of the top ranked docked pose of complex B revealed that NTD (N-terminal domain) of spike protein interacted with the CRD domain of DC-SIGN (Figure 7B & Table 2). Since it was known that Spike protein interacted with SP-D through receptor binding domain (RBD) (24), we postulated that Spike protein could interact with both SP-D and DC-SIGN (CRD) through two distinct RBD and NTD domains, respectively. This inference is further supported by the *in vitro* observation that binding of DC-SIGN and Spike protein was enhanced by rfhSP-D (Figure 1C). Tripartite complex was generated by docking complex A (DC-SIGN and SP-D) with Spike protein. The top two docked poses (complexes C1 and C2) were analysed for intermolecular interactions (Figure 8; Table 2). In both C1 (Figure 8A) and C2 (Figure 8B) complexes, DC-SIGN (CRD) interacted with NTD domain of Spike protein. In C1, there were no molecular interactions between Spike protein and rfhSP-D (Figure 8A; Table 2). In C2, Spike protein interacted with rfhSP-D through RBD (Figure 8B; Table 2).

**Table 2:**
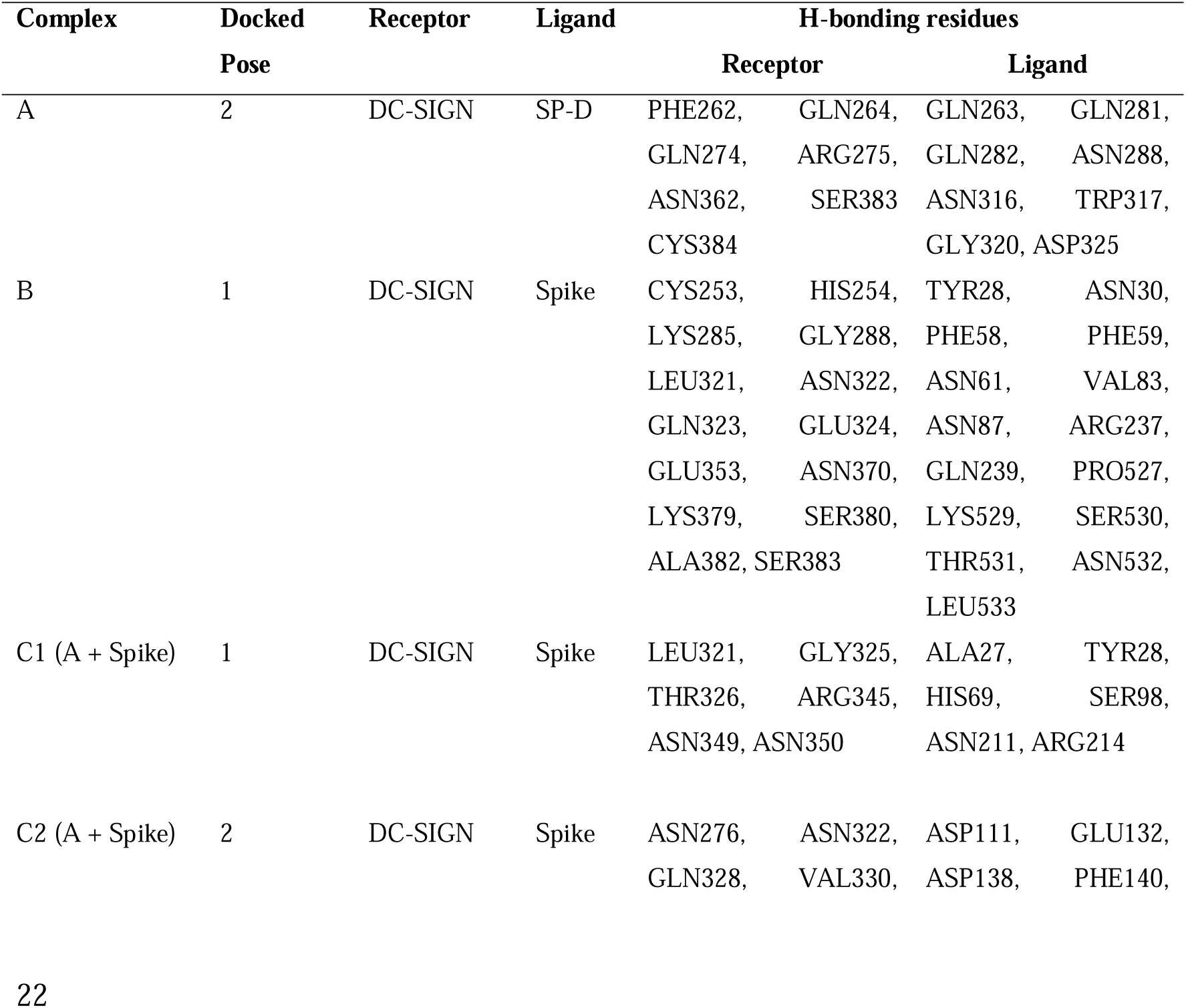
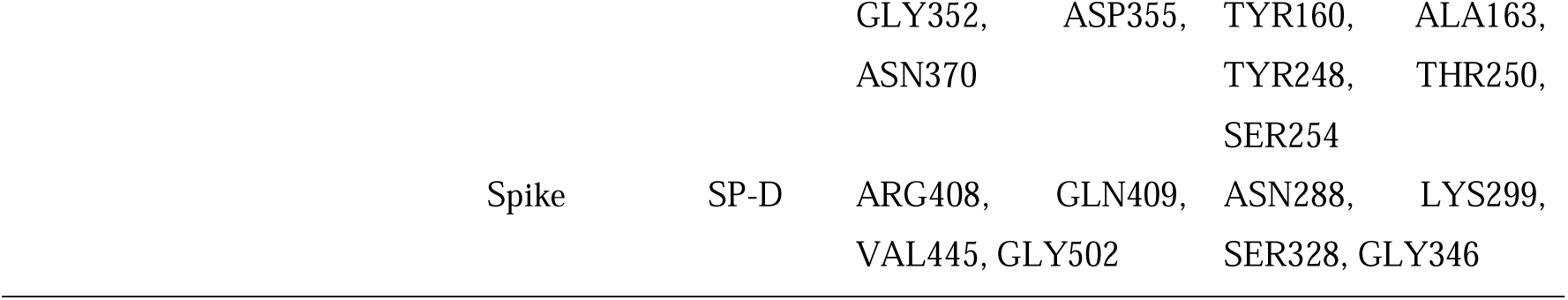
Interaction analysis of the docked complexes of DC-SIGN, spike and SP-D

**Figure 7:**
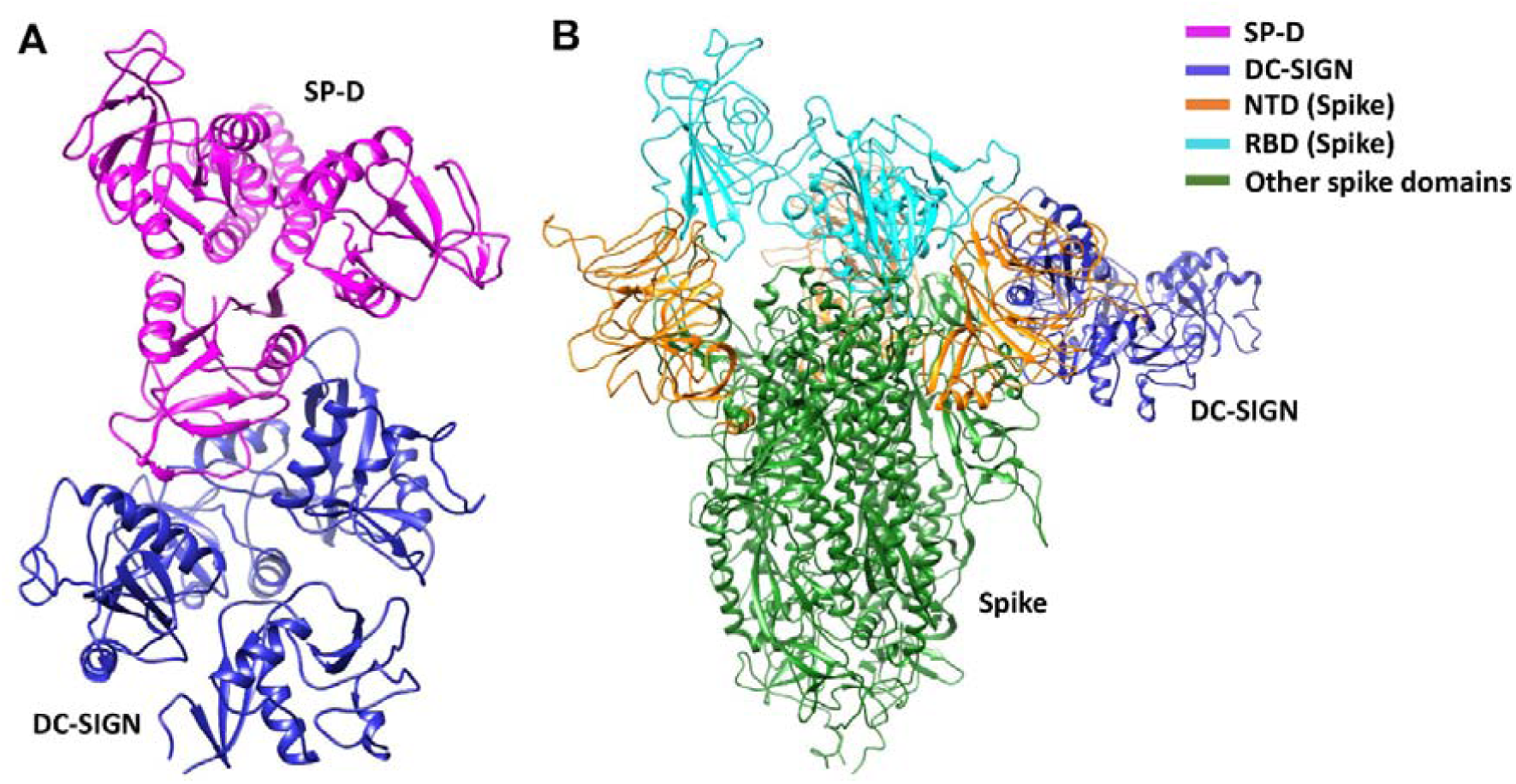
DC-SIGN interacts with both SP-D and SARS-CoV-2 spike. Docked poses of (A) complex A, and (B) complex B selected for docking and MD simulations respectively. In complex B, spike interacts with DC-SIGN(CRD) through the NTD domain (orange).

**Figure 8:**
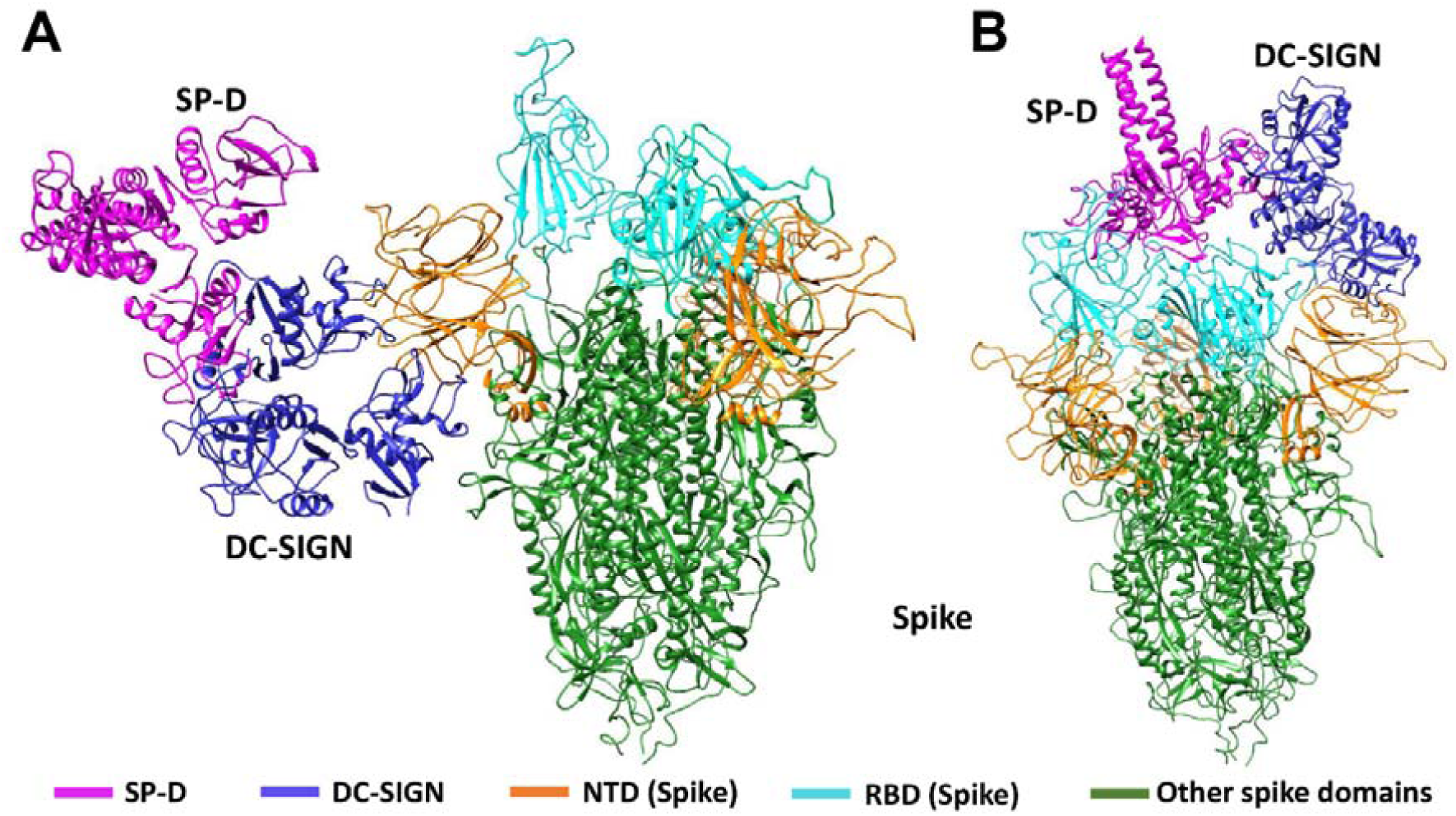
Tripartite complex of SP-D, DC-SIGN and SARS-CoV-2 Spike. Docked poses of tripartite complexes selected for MD simulation analysis. In complex C1, DC-SIGN (CRD) interacts with NTD of spike (A); and in complex C2, DC-SIGN (CRD) interacts with NTD of spike and SP-D interacts with RBD of spike (B).

### SP-D Stabilises DC-SIGN and SARS-CoV-2 Spike protein Interaction

MD simulations were performed to assess the effect of SP-D on DC-SIGN (CRD) and Spike protein interaction. The root mean square deviation (RMSD) of complexes C1 and C2 was lesser than complex B through the course of simulation, indicating that the binding of SP-D enhances the stability of DC-SIGN and spike interaction (Figure 9A). This observation was supported by potential energy (PE), distance, and H-bond profile. Trajectory analysis of PE, intermolecular distance and H-bonds between DC-SIGN and spike indicated higher stability of C1 and C2 complexes as compared to B (Figures 9B, 10A-C, 10D-F). Between the tripartite complexes, C1 exhibited slightly better stability than C2 (Figures 9 & 10). These analyses suggest that the interaction of DC-SIGN and spike gets stabilized in the presence of SP-D.

**Figure 9:**
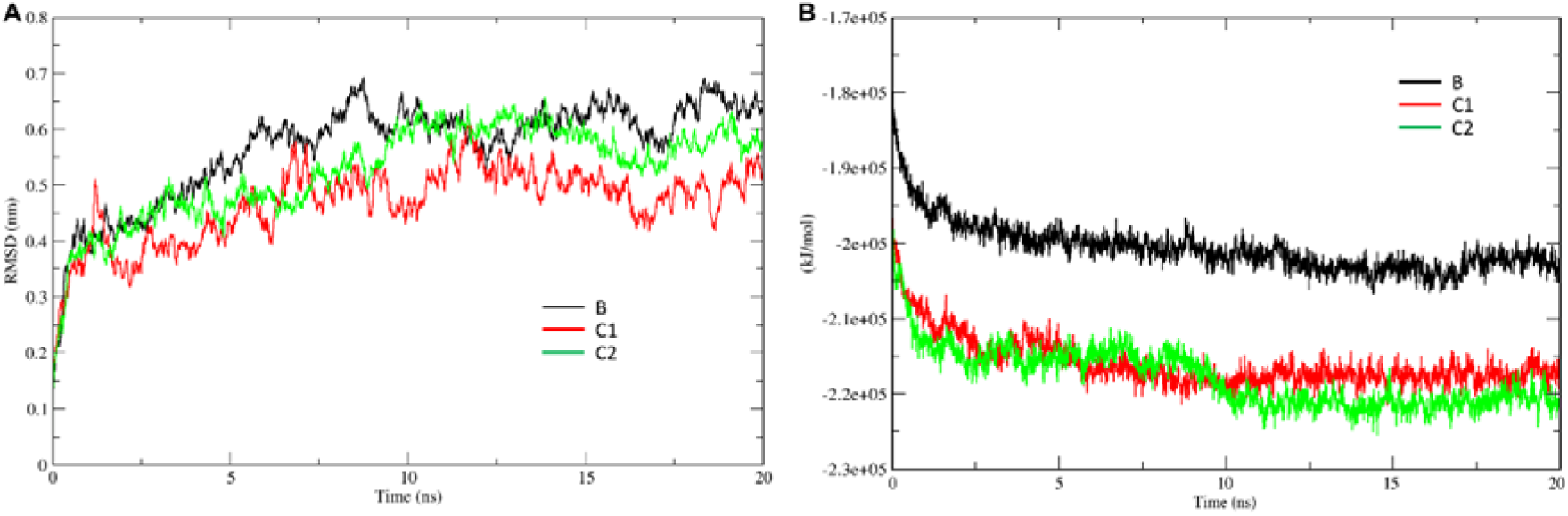
SP-D stabilises SARS-CoV-2 Spike interaction with DC-SIGN. Comparative MD simulation profile for complexes B, C1 and C2 of (A) root mean square deviation (RMSD) and (B) potential energy (PE). RMSD and PE of C1 and C2 are lesser than B indicating stability of tripartite complexes.

**Figure 10:**
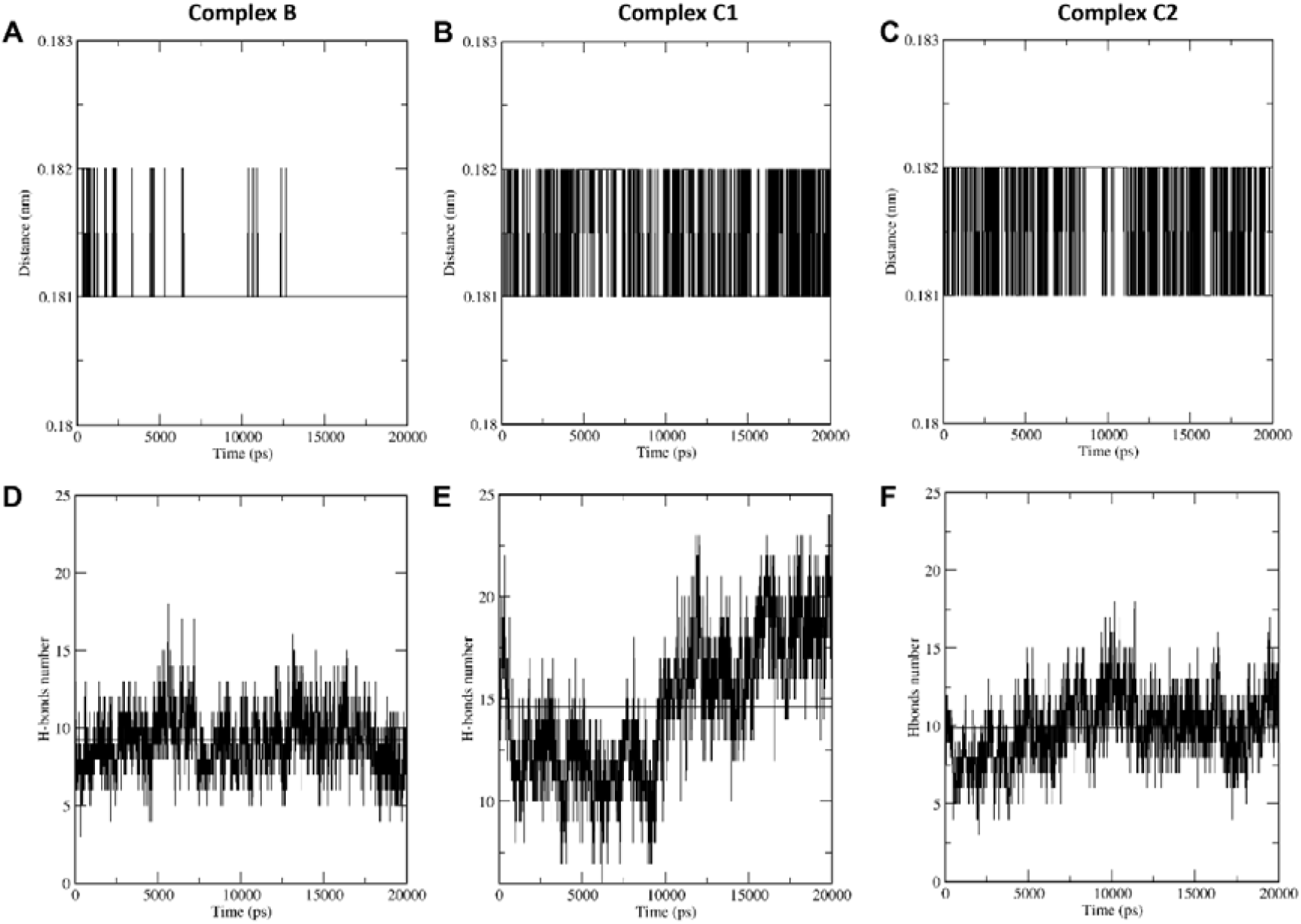
SP-D stabilises SARS-CoV-2 Spike interaction with DC-SIGN. Comparative MD simulation profile of complexes B, C1 and C2 for average distance (A, B & C) and H-bonds (D, E & F) between DC-SIGN and spike. Its observed that the intermolecular distance is conserved across the simulation period for tripartite complexes C1 and C2 as compared to complex B. The number of intermolecular H-bonds between DC-SIGN and spike are also higher for complexes C1 and C2 as compared to B. These observations indicate the stabilising effect of SP-D on spike and DC-SIGN(CRD) interaction.

## Discussion

Specific molecular structures on the surfaces of pathogens (PAMP) are directly recognised by pattern recognition receptors (PRRs) (4). PRRs serve as a link between nonspecific and specific immunity. PRRs can exert nonspecific anti-infection, anti-tumour, and other immune-protective actions by recognising and binding ligands (46). CLRs belong to PRRs, which use calcium to recognise carbohydrate residues on harmful bacteria and viruses (47). DC-SIGN and SP-D are examples of CLRs that play an important role in anti-viral immunity, including SARS-CoV-2, the causative agent of coronavirus induced illness 2019 (COVID-19) (48, 49). The availability of virus receptors and entry cofactors on the surface of host cells determines tissue tropism for many viruses (50, 51). We found rfhSP-D potentiated SARS-CoV-2 binding and entry to DC-SIGN expressing cells. Furthermore, rfhSP-D treatment was also found to impair downstream signalling induced by the binding of Spike protein to DC-SIGN resulting in the downregulation of pro-inflammatory mediators′ gene expression. However, further experiments using DC-SIGN expressing cells challenged with rfhSP-D treated SARS-CoV-2 clinical isolates need to be undertaken to confirm the rfhSP-D mediated cytokine modulation observed. Our findings elucidate a novel interaction between rfhSP-D, DC-SIGN and SARS-CoV-2, which may uncover therapeutic potential for controlling SARS-CoV-2 infection and subsequent cytokine storm.

DC-SIGN, widely expressed on DCs and alveolar macrophages in the lungs, interacts with SARS-CoV-2 through Spike protein (52). DC-SIGN has previously been shown to interact with SARS-CoV, HIV-1 and Ebola virus (41, 53, 54). SARS-CoV uses DC-SIGN for entry into DCs (52). In addition to its role as a virus attachment receptor, DC-SIGN has been implicated in triggering DC maturation, myeloid cell cytokine response, and T cell priming. Another CLR, SP-D, has been shown to have antiviral properties against SARS-CoV-2, HIV-1 and IAV infection (55, 56). We previously demonstrated that rfhSP-D reduced SARS-CoV-2 S1 protein binding to HEK293T cells overexpressing ACE2 receptors and infection in A549 cells by restricting viral entry (25). However, the role of SP-D in SARS-CoV-2 and DC-SIGN interaction is not well understood.

The binding of the SARS-CoV-2 Spike protein to the host cell via the ACE2 receptor is one of the critical steps in the SARS-CoV-2 infection (57). The receptor binding motif (RBM) (455-508) within the RBD of S1 protein interacts with the virus-binding residues consisting of Lys31, Glu35, and Lys353 of dimeric ACE2 (58). Although the sequence of events around the Spike protein/ACE2 association is becoming more evident, additional factors that aid infection remains unknown, for example, SARS-CoV-2 transport to the ACE2 receptor (36). Both SARS-CoV and SARS-CoV-2 Spike proteins have the same affinity for ACE2, but the transmission rate are drastically different (37). It has been suggested that the higher transmission rate of SARS-CoV-2 compared to SARS-CoV is due to more efficient viral adherence via host-cell attachment factors, leading to more efficient infection of ACE2 expressing cells (38, 39). DC-SIGN has also been identified as a SARS-CoV Spike protein receptor capable of enhancing cell entry in ACE2^+^ pneumocytes via DC transfer (40). Recently, it has been shown that DC-SIGN binds to SARS-CoV-2 Spike protein and promotes trans-infection (20). In this study, we investigated the potential of rfhSP-D in inhibiting SARS-CoV-2 binding and entry into DC-SIGN expressing cells. Targeting viral entry into a host cell is a new technique for creating and developing antiviral medicines that stop viral propagation early in the SARS-CoV-2 viral cycle (59). We have independently confirmed the previously reported protein interactions between SARS-CoV-2 and rfhSP-D or DC-SIGN (20, 24, 25).

Here, we show that rfhSP-D enhances the binding of the SARS-CoV-2 Spike to DC-SIGN. This is further confirmed by *in-silico* molecular dynamics studies which indicate that SP-D stabilises the binding interactions between DC-SIGN CRD and N-terminal domain of SARS-CoV-2 Spike protein. The consequence of this tripartite complex involving DC-SIGN, SARS-CoV-2 Spike protein and rfhSP-D on viral infection was assessed using SARS-CoV-2 Spike protein-expressing replication-incompetent lentiviral pseudotyped viral particles since they are a safe alternative to the live virus. Using these pseudotypes, we demonstrate that rfhSP-D enhances spike protein binding and uptake in DC-SIGN expressing cells. A significant increase in spike protein binding and transduction was observed compared to untreated samples (Cells + SARS-CoV-2) to rfhSP-D (20 µg/ml) treatment. It has been shown previously that SP-D enhances the clearance of IAV from the lung *in vivo* (60). Similarly, the interaction of rfhSP-D with DC-SIGN may augment SARS-CoV-2 binding and uptake by macrophages, indicating that rfhSP-D may promote the clearance of SARS-CoV-2 via DC-SIGN.

The effect of rfhSP-D on gene expression levels of pro-inflammatory mediators in SARS-CoV-2 Spike protein challenged DC-HEK and DC-THP-1 cells were investigated in the current study. To our knowledge, this is the first study looking at the impact of rfhSP-D on DC-SIGN cells challenged with SARS-CoV-2. rfhSP-D showed anti-inflammatory effects on DC-SIGN expressing cells, as evident from the reduction in the levels of cytokines/chemokines such as TNF-α and IL-8.

DC-SIGN present on DC surface has been implicated in activating the STAT3 pathway during viral infection (61, 62). STAT3 plays a crucial role in activating transcription factor NF-κB in SARS-CoV-2 infection in myeloid cells, which may trigger subsequent cytokine production and stimulate pathological inflammation (63, 64). The activation of NF-κB in viral infection induces gene expression of a wide range of cytokines (e.g., IL-1, IL-2, IL-6, IL-12, TNF-α, LT-α, LT-β, and GM-CSF), and chemokines (e.g., IL-8, MIP-1, MCP1, RANTES, and eotaxin) (65). These inflammatory mediators are involved in antiviral immunity and essential for infection resistance (65). Nevertheless, in moderate and severe SARS-CoV 2 infection, the activation of NF-κB in various cells, including macrophages in the lungs, liver, kidney, central nervous system, gastrointestinal system, and cardiovascular system, results in the production of IL-1β, IL-6, IL-8, and TNF-α (66). This may result in cytokines storm and organ failure and, consequently, morbidity and mortality (66, 67). Immunomodulation at the level of NF-κB activation and inhibitors of NF-κB degradation may reduce the cytokine storm and lessen the severity of SARS-CoV-2 infection (66, 68). Pro-inflammatory mediators have been shown to be induced by SARS-CoV-2 Spike protein in THP-1 cells as in vitro model for lung macrophages (69). In this study, the inflammatory response was evaluated via measuring the gene expression levels of NF-κB in DC HEK and DC THP-1 challenged with SARS-CoV 2 spike protein. Our findings show rfhSP-D downregulates the gene expression levels of NF-κB in DC-HEK and DC-THP-1 challenged with SARS-CoV-2 Spike protein compared with the control. Thus, rfhSP-D suppresses pro-inflammatory immune response in DC-SIGN expressing immune cells.

Another critical element in the pathophysiology of SARS-CoV-2 infection is TNF-α, which is produced in the airway by macrophages, mast cells, T cells, epithelial cells, and smooth muscle cells (4, 70). TNF-α synthesis is predominantly stimulated by PAMPs through NF-κB activation (71). Various studies have reported that patients with severe SARS-CoV-2 infection display elevated plasma levels of TNF-α (72-74). This causes airway inflammation due to recruitment of mostly neutrophils (75). In addition, TNF-α stimulates the production of cytokines like IL-1β and IL-6 (76). DC-HEK and DC-THP-1, challenged with SARS-CoV-2 Spike protein, pre-treated with rfhSP-D, caused downregulation in the gene expression levels of TNF-α as compared to rfhSP-D-untreated cells. The results suggested an important immunomodulatory role of rfhSP-D in SARS-CoV-2-mediated inflammation.

IL-1β is released after activating the inflammasome in response to a variety of infections, including viruses such as SARS-CoV-2 (77). When compared to non-infected subjects, high levels of IL-1β were found in the plasma of severe as well as moderate CoVID-19 cases (73). Cell pyroptosis is a highly inflammatory form of programmed cell death typically seen with cytopathic viruses. An increase in IL-1β production is a downstream sign of pyroptosis (78). As a result, pyroptosis plays an essential role in the pathogenesis of SARS-CoV-2 and is a likely trigger for the uncontrolled inflammatory response (79). In this study, DC-THP-1 cells challenged by Spike protein, pre-treated with rfhSP-D, exhibited low mRNA levels of IL-1β as compared to the control. Thus, rfhSP-D may reduce the unnecessary inflammatory response to SARS-CoV-2 infection via reduction in IL-1β production.

IL-6 is a glycoprotein that regulates the immune system, haematopoiesis, inflammation and is a major player in SARS-CoV-2 infection (80). Many cell types, including T and B lymphocytes, monocytes/macrophages, dendritic cells, fibroblasts, and endothelial, express IL-6 (80, 81). A higher level of IL-6 in the plasma has been linked with the severity of SARS-CoV-2 infection (82, 83). rfhSP-D-treated DC-THP-1 cells challenged by Spike protein showed downregulation of IL-6 transcripts as compared with untreated cells. This suggests a role for SP-D in preventing IL-6 immunopathogenesis due to SARS-CoV-2.

IFN-α is a cytokine mainly secreted by virus-infected cells associated with stimulation of immune response and limiting viral infection (84). Nonetheless, elevated expression levels of IFN-stimulated genes (ISGs) have been triggered by SARS-CoV 2, which exhibits immunopathogenic potential (58). IFN-α expression levels are downregulated in rfhSP-D treated DC HEK cells challenged with SARS-CoV 2 spike protein compared to the control. The results suggest rfhSP-D may elevate immunopathology potential of SARS-CoV-2.

Another element that may aid viral infectivity of DCs is MHC class II molecule. SARS-CoV-2 has been shown to upregulate MHC class II gene expression (85). High expression level of MHC class II molecules on the surface of antigen-presenting cells is crucial for regulating and inducing an adaptive immune response to respiratory viruses (86). However, limited- expression levels of MHC class II molecules on type II alveolar cells and macrophages improve respiratory viral disease outcomes (87). The binding of SARS-CoV-2 Spike protein to THP-1 cells polarises towards M1-like phenotype together with an increase in MHC class II molecules (69). DC-THP-1 cells challenged with SARS-CoV-2 Spike protein and treated with rfhSP-D showed downregulation of MHC class II mRNA expression levels. Thus, SP-D may have a role in modulating antigen presentation in order to avoid an unwanted and exaggerated adaptive immune response.

Chemokines, such as IL-8 and RANTES, are vital for recruiting inflammatory cells from the intravascular space across the endothelium and epithelium to the inflammation site (88). IL- 8, commonly known as CXCL8, is a crucial mediator of inflammation with a direct chemotactic and priming action on neutrophils (89). In addition, IL-8 induces NETosis (Neutrophil extracellular traps/NETs). SARS-CoV-2 infected patients exhibit elevated levels of citrullinated histone H3 (Cit-H3) and myeloperoxidase (MPO)-DNA, which are specific markers of NETs that may cause organ damage (90). DC-THP-1 challenged with Spike protein and treated with rfhSP-D had low mRNA levels of IL-8 as compared to the control. There appears a role for SP-D in preventing IL-8-associated pathogenies due to SARS-CoV-2.

RANTES (CCL5) is a chemokine that has been linked with enhanced pathogenicity and mortality in SARS-CoV-2 infection (91). Compared to healthy control, SARS-CoV-2 infected patients contain higher serum RANTES and IL-6 levels which correlated with severity of CoVID-19 (92). In this study, the mRNA expression levels of RANTES were found to be considerably downregulated in DC-HEK cells challenged with Spike protein and treated with rfhSP-D. Thus, SP-D may modulate leukocyte recruitment to infection areas.

Our study reveals that rfhSP-D can effectively increase the binding and uptake of SARS-CoV-2 by DC-SIGN expressing cells. We also show that rfhSP-D exhibit substantial effectiveness in downregulating virus-induced inflammatory response in DC-SIGN expressing cells. However, further study is required to assess the expression of DC-SIGN in individuals with mild/severe SARS-CoV-2 infection. SARS-CoV-2 is also known to affect organs other than the lungs. Thus, it is imperative to study the possibility of viral transfer to secondary sites via DC-SIGN and the effect of SP-D on this process. Additionally, the effects of rfhSP-D mediated DC-SIGN: SARS-CoV-2 interaction needs to be studied in the lung microenvironment using established animal models for CoVID-19 such as Hamsters, Mouse, Ferret, Mink, Tree Shrew, and Non-human Primates. In conclusion, our data suggests that rfhSP-D stabilises the interaction between SARS-CoV-2 Spike protein and DC-SIGN and helps in increasing viral uptake by macrophages, suggesting an additional role for rfhSP-D in SARS-CoV-2 infection.

## Funding

CK and SI-T are grateful to the grants received from the Department of Biotechnology, India [No. BT/PR40165/BTIS/137/12/2021]. NT and MMN are funded by the Wellcome Trust (GB-CHC-210183)

## Notes

### Competing Interest Statement

The authors have declared no competing interest.

